# Inhibition of Proteasome Activity Facilitates Definitive Endodermal Specification of pluripotent Stem Cells by influencing YAP signaling

**DOI:** 10.1101/2024.03.15.585134

**Authors:** Akshaya Ashok, Ashwini Ashwathnarayan, Smitha Bhaskar, Spandana Shekar, Jyothi Prasanna, Anujith Kumar

## Abstract

Understanding the molecular players that control the specification of definitive endoderm is imperative to obtain the homogenous population of pancreatic β-cells from stem cells. Though the Ubiquitin proteasome system (UPS) has been envisaged as a crucial intracellular protein degradation system, its role in germ layer specification remains elusive. In this study, using a mouse embryonic stem cells model system (mESCs) we observed decreased proteasomal activity specifically during endoderm, but not in meso- or ecto-derm differentiation. Extraneous inhibition of proteasomal activity during differentiation enhanced the expression of endodermal genes specifically. Enhancing proteasomal activity by including the activator IU1 in the induction culture, inhibited definitive endodermal differentiation. Further, inhibiting proteasomal activity at the definitive endodermal stage resulted in enhanced generation of insulin-positive cells. A similar increase in endodermal gene expression by inhibiting proteasomal activity was observed in miPSC and hiPSC differentiated towards endodermal lineage. Mechanistic insight showed no contribution of endoplasmic reticulum unfolded protein response but revealed the involvement of the YAP signaling pathway in proteasome-inhibited enhanced endodermal differentiation. Unravelling the specific involvement of UPS in endodermal cell fate specification in pluripotent stem cells paves the way for obtaining better qualitative and quantitative definitive endodermal cells for plausible cellular therapy in the future.

## Introduction

The worldwide increasing prevalence of Diabetes mellitus (DM) has been a major public health problem approaching an epidemic proportion (1). Conventional therapeutic strategies for this condition like drug regimens and insulin injections are increasing the risk factors to the patients and result in severe diabetes-associated diseases such as kidney disease, peripheral neuropathy, coronary heart failure, etc (2, 3). These limitations have compelled us to glean the current hypothetical alternative approach of stem cell-derived β-cell transplantation (4, 5). However, to execute the approach, an in-depth understanding of the cascade of signalling pathways that regulate pancreatic β-cells development is imperative.

The major dictator of pancreatic development relies on the stage-dependent synchronized expression of different combinations of transcription factors (TFs) (6, 7). The homeostasis of these TFs is majorly determined by the balance between transcription and post-transnational regulation via the ubiquitin proteasome system (UPS) (8, 9). The UPS is a major protein degradation system that targets 70-80% of the intercellular proteins (10). Proteasomes cleave most of the intracellular proteins into oligopeptides and these small peptides are further cleaved by peptidases (oligopeptidases or aminopeptidases) to release free amino acids that are further utilized for protein synthesis (11). Protein substrates targeted for degradation are polyubiquitinated via covalent bond by sequential activation of Ubiquitin-activating enzyme (E1) which activates and transfers the ubiquitin to a carrier enzyme E2 and finally, E3 (ubiquitin-protein ligase) enzyme ubiquitinates the target substrate that is relayed to the proteasome for degradation (12).

Several recent studies have accentuated the key role played by UPS in stemness and differentiation of stem cells to different lineages. Enhanced Proteasomal activity was observed in pluripotent hESCs compared to their differentiated counterpart and inhibition of proteasomal activity promoted the differentiation (13, 14). In pancreatic β-cells, the importance of UPS in maintaining protein homeostasis has been well demonstrated (15). For instance, the degradation of homeodomain transcription factor (TF) PDX1 occurs by GSK3-dependent phosphorylation of serine residues in PDX1, which acts as a signal for ubiquitination and proteasomal degradation (16). The increased degradation of PDX1 results in β-cell dysfunction and culminates in diabetic conditions (17). *In vivo* studies have shown the PDX1 C-terminus–interacting factor-1 (Pcif1) targets PDX1 for ubiquitination and proteasomal degradation (8). Another important transcription factor, MafA, a glucose-regulated pancreatic β-cell-specific transcriptional activator that regulates insulin gene expression has also been shown to be the substrate of the proteasome (18, 19). Phosphorylation of MafA at multiple serine and threonine residues by GSK3 and MAP kinases mediates its degradation (20). Insulin secretion by β-cells has been shown to be regulated by UPS. Inhibition of proteasome activity decreased the VDCC calcium channel and in turn decreased the insulin secretion in isolated mouse pancreatic islets (21). During the exposure to different concentrations of glucose, the expression of genes in the UPS pathway is also modulated and the blockade of proteasome degradation machinery enhanced glucose-stimulated insulin secretion (22). All these studies implied the importance of UPS in the proper functioning of matured β-cells. However, no information is available that highlights the role of UPS during the β-cell differentiation from stem cells.

In the present study, we have demonstrated the unconventional role of UPS in β-cell generation and found that inhibition of proteasomal activity favours the endodermal lineage formation as compared to other lineages during targeted differentiation of mouse and human pluripotent stem cells. Further, an in-depth investigation of the mechanism showed the crosstalk between UPS and the Hippo-Yap pathway to be responsible for the enhanced endodermal differentiation.

## Experimental Procedure

### Cell culture

The R1 mESCs and PDX1-GFP iPSCs (expression of GFP driven by PDX1 promoter) were cultured on mitomycin-inactivated mouse embryonic fibroblast (MEF) with ESC medium [DMEM High Glucose medium (Gibco-#11965-092) with 15% FBS (Gibco-#10270-106), and supplement of 2mM L-Glutamine (Gibco-#11539-876), 1X Sodium pyruvate (Gibco-#11360-070), 1X Penicillin/streptomycin (Gibco-#15240-062), 1X non-essential amino acids (Gibco-#11140-050), 100µM β-mercaptoethanol (HiMedia-#MB041) and 100units/mL leukaemia inhibitory factor (LIF) (Millipore-#ESG1106). The hiPSCs (human pluripotent stem cells) were cultured on 1% matrigel (Corning-#354277) with mTESR medium (Stem Cell-#100-0274). 10 µM Y27632 (Tocris-#1254) was added on the day of cell seeding and after 24 hrs, the media was replaced with mTESR lacking Y27632.

### Embryoid body (EB) formation

mESCs were allowed to form embryoid bodies in the low attachment 6 well plates (Corning-#3471) in ESC medium lacking LIF. The medium was changed on every alternate day for 14 days and where required cells were cultured in the presence or absence of MG132 (Proteasome inhibitor-PI) (Sigma-#C2211) and Proteasome activator (IU 1-IU) (Sigma-#I1911).

### Mouse pluripotent stem cell β-cell Differentiation

The protocol was adapted from a previously published study with minor modifications (23). mESCs were split and approximately 6 × 10^4^ cells were plated onto a 12-well plate coated with 1% matrigel in ESCs medium without LIF and the next day medium was replenished with differentiation medium containing 40% MCDB (Sigma-#M6770), 100µM β-mercaptoethanol, 100nM Vitamin C (Sigma-#A4403), 1X ITS & 1X LABSA (Sigma-#I3146&#L9530) and 1X Penicillin/streptomycin and DMEM Low Glucose medium (Gibco-#11885-084). The following combinations of cytokines were added at different stages of differentiation. Day 1: mESCs were cultured in a differentiation medium containing 75ng/mL Activin (Sino Biological-#10429), 3μM/mL CHI99021 (Tocris-#4423) and 4% FBS. Day 3: Medium was changed with differentiation medium containing activin and 3% FBS. Day 6: The medium was changed with the differentiation medium containing 0.25μM Sant-I (Tocris-#1974), 100ng/mL Noggin (Sino Biological-#10267), 2μM Retinoic acid (RA) (Sigma-#R2625), 50ng/mL KGF (Peprotech-#100-19) and 2% FBS. Day 8: A half medium change was given with the differentiation medium containing 0.25μM Sant-I, 2μM RA, 50ng/mL KGF and 1% FBS. Day 10: A complete medium change was given with a differentiation medium containing 50ng/mL KGF, 10μg Heparan sulphate (Sigma-#H7640) and 1% FBS. Day 16: Medium change was given with differentiation medium containing 50ng/mL β-cellulin (Peprotech-#100-50), 10nM Exendin (Prospec-#HOR-269), 10mM Nicotinamide (Sigma-#N0636) and 50ng/mL GDF II (Peprotech-#120-11). Cells were harvested on day 22 for matured β-cell transcript and protein expression analysis.

### hiPSC Endoderm Differentiation

Differentiation protocol was adapted from Pagliuca et al. (24) with minor changes. Approximately, 0.5×10^6^ cells were plated on matrigel-coated 12 well plates with 10µM Y27632 in mTESR medium overnight and the medium was replenished with fresh mTESR medium and cultured for 24-48 hours. Day 1: The S1 ifferentiation medium [44mg Glucose (Sigma-#G8644), 246mg NaHCO_3_ (Sigma-#S5761), 2g BSA (HiMedia-#TC548), 2µL ITS (Sigma-#I3146), 1mL Glutamax (Gibco-#35050-061), 4.4mg Vitamin C (Sigma-#A4403), 1X Penicillin/streptomycin (Gibco-#15240-062), make up to 100mL with MCDB (Sigma-#M6770)] with 100ng/mL Activin (Sino Biological-#10429) and 14μg/mL CHI99021 (Tocris-#4423). Day 2: The medium was changed to S1 differentiation media with 100ng/mL Activin. Day 4: The S2 differentiation medium [44mg Glucose, 123mg NaHCO_3_, 2g BSA, 2µL ITS, 1mL Glutamax, 4.4mg Vitamin C, 1X Penicillin/streptomycin, make up to 100mL with MCDB] with 50ng/mL KGF (Peprotech-#100-19). Day 7: Medium was changed to S3 differentiation medium [44mg Glucose, 123mg NaHCO_3_, 2g BSA, 500µL ITS, 1mL Glutamax, 4.4mg Vitamin C, 1X Penicillin/streptomycin, make up to 100mL with MCDB] with 100ng/mL Noggin (Sino Biological-#10267), 50ng/mL KGF, 0.25μM Sant-I (Tocris-#1974), 2μM Retinoic Acid (Sigma-#R2625), 500nM PdbU (Cell signalling Technology-#12808) and 10μM Y27632. Day 8: Medium was changed to the same condition as Day 7 without the Noggin supplement. Day9: The S3 differentiation medium with 5ng/mL Activin, 50ng/mL KGF, 0.1μM RA, 500nM PdbU, 10μM Y27632 and 0.25μM Sant-I. Day11: Medium was changed to S3 differentiation medium with 5ng/mL Activin, 50ng/mL KGF, 0.25μM Sant-I, 0.1μM RA, 500nM PdbU and 10μM Y27632. Cells are harvested for transcript and protein expression analysis on each day mentioned above.

### Mesodermal Differentiation

Approx. 1×10^6^ mESCs/sq.cm plated with KnockOut DMEM (Gibco-10829-018) supplemented with 2% FBS and incubated overnight at 37°C and 5% CO_2_. The medium was changed to differentiation media supplemented with 10 ng/mL each of bFGF (Peprotech-#100-18B), PDGF (R&D System-#220BB) and EGF (Abclonal-#RP01030) with every alternative day medium change. The cells were harvested on day 6 for transcript and protein expression analysis.

### Ectodermal Differentiation

mESCs at a density of 2×10^5^ cells/sq.cm were cultured in ESC medium overnight and changed to neural differentiation medium containing DMEM F12 (Himedia-#AL139A), 1X N2 & B27 supplement (Gibco-#17502-048&#12587-010) at a 1:1 ratio with 10^-8^M Retinoic acid. The cells are harvested on day 6 for transcript and protein analyses.

### Proteasomal activity assay

Cells were homogenized with homogenization buffer [50mM Tris Hcl pH 7.5, 250mM Sucorse, 5mM MgCl_2_,0.5mM EDTA, 2mM ATP, 1mM DTT, 0.025% Saponin] (25) using a probe sonicator (Misonix). Homogenized samples were centrifuged at 12000g for 15min at 4°C and the supernatant protein was collected, and protein concentration was estimated by BCA Kit (Takara-#T9300A). For the assay, 10-20µg of protein lysate and 200µM proteasome-specific substrate Suc-LLVY-AMC (Enzo-#BML-P802-0005) were added and the reaction mix was made up to 100µL with activity buffer [50mM Tris Hcl pH 7.5, 40mM KCl, 5mM MgCl_2_,0.5mM ATP, 1mM DTT] and the reaction mixture incubated at 37°C for 1 hour. The reaction was arrested with the addition of 400µL of 100% ethanol (HiMedia-MB106) and the absorbance of the samples was measured at the wavelength of 370-430nm excitation and emission respectively in Ensight multimode plate reader (Perkin Elmer).

### Reverse Transcriptase PCR and Real-Time PCR

RNA was extracted by using RNAiso Plus (Takara-#9109) according to the manufacturer’s instruction and then quantified by Nanodrop One at 260nm (Thermo Scientific). Complementary DNA (cDNA) synthesis was carried out with 1 μg of RNA using the PrimeScript RT reagent kit (Takara-#RR037A-4). The cDNA was diluted and semiquantitative and Real-time PCR techniques were performed. semiquantitative PCR was performed using 2X EmeraldAmp GT PCR Master Mix (Takara-#RR310) and gene-specific primers and the amplified PCR products were visualized on 2% agarose gel. qPCR was performed by using TB Green master mix (Takara-#RR820) using gene-specific primers in a 7500 Applied Biosystems Real-time PCR machine. The PCR primers designed encompassing the introns are tabulated in Supplementary Table 1.

### Immunofluorescence staining

Cells cultured in 12/24well plates were fixed with 4% PFA (Sigma-#HT5014) for 20 min. at room temperature. After PBS washes cells were permeabilized with 0.5% Triton X-100 (Sigma-#X100RS) for 20 min and then blocked with 20% FBS. Cells were overnight incubated with 1:400 concentration of primary antibody diluted with PBS at 4°C. The following primary antibodies were used for immunofluorescence staining: Mouse anti-Ki-67 (BD Bioscience-#550609), Rabbit anti-Hnf6 (Abclonal-#A12774), and Rabbit anti-INS (Cloudclone-#PAA448Hu01). After three washes with the PBS, corresponding secondary antibodies (AlexaFluor 594 Goat Anti-Mouse (Invitrogen-#A11005, AlexaFluor 594 Donkey Anti-Rabbit (Invitrogen-#A21207) with 1:800 dilution with PBS were incubated for 2 hours at room temperature and the nuclei were stained with DAPI (Sigma-#D9542). Cells were rinsed with fresh PBS and fluorescence-stained images were captured by using Q-capture software on an Olympus 1X73 inverted Fluorescence microscope and the images were processed with the Image J software.

### Western blotting

Cells were lysed in RIPA lysis buffer and the clear protein supernatant was collected after centrifugation at 10000 rpm at 4°C for 20-30min. 25-30µg protein was loaded on resolving 12% SDS-PAGE and later transferred to a methanol-activated PVDF membrane (Millipore-# IPVH00010) by using a semi-dry Trans blot apparatus (Bio-Rad). The membrane was blocked with 5% BSA (Sigma-#A1470)/ 5% Skimmed milk in 1X TBST and incubated with primary antibodies at 1: 1000 dilution overnight at 4°C on a rocker. The primary antibodies used were Mouse anti-beta actin (Abclonal-#AC004), Rabbit anti-Foxg1 (Abcam-#ab18259), Rabbit anti-Cxcr4 (Cusabio-#PA006254YA01Hu), Rabbit anti-Vimentin (Cusabio-#PA025857LA01Hu), Rabbit anti-Pdx1 (Abclonal-#A10173), Rabbit anti-Ubiquitin (Cell signalling Technology-#3933S) and Rabbit anti-Yap (Abclonal-#A1002)]. Post three washes with the 1X TBST, the membrane was incubated with appropriate HRP conjugated secondary antibodies (Abcam-Goat Anti Mouse IgG #ab97023, Goat Anti Rabbit IgG #ab97051) at a 1:2000 concentration for 1 hour 30 min. at room temperature and finally, blots were developed by using ECL HRP substrate (Advansta-#K12045-D20) on Chemi Doc XRS+ system (BioRad).

### Flow cytometry

Cells were trypsinized and washed with cold 1X PBS (Takara-#T9181) and fixed with 0.5% PFA for 30 min. at room temperature. Cells were washed with 1X PBS and were permeabilized with cold 1X Permwash (BD Biosciences-#554723) for 30 min. and blocked with cold 20% FBS in 1X Permwash for 30 min. at room temperature on a rocker. Further cells were incubated with primary antibody at a1:200 dilution at 4°C on a rocker platform for overnight. Post three washes with the 1X PBS, respective secondary antibodies were added at a concentration of 1:800 for 1 hour 30 min. at room temperature and cells were washed with 1X PBS. At least 10,000 events were acquired on the FACS Calibur cytometer (BD Bioscience) and analyzed by FACS Diva software. GFP expression in the differentiated Pdx1-GFP iPSCs was analyzed in trypsin-dissociated live single cells, resuspended 1X PBS and acquired the GFP positive cells on FACS Calibur.

### TPE-MI Staining for unfolded protein quantification

Cells were rinsed with PBS treated with 50µM TPE-MI (MedChemExpress-#HY-143218) and incubated for 30min, at 37°C in a CO_2_ incubator. Cells were harvested in the RIPA lysis buffer and the clear protein supernatant was separated by centrifugation at 10K rpm. The supernatant protein concentration was determined by BCA kit and nearly 20µg protein from each condition was used to measure the unfolded protein load. Then, TPE-MI fluorescence was recorded at 350-470nm (Excitation/emission) by using an Ensight multimode plate reader (Perkin Elmer).

### Statistical analysis

Statistical significance was analyzed by a tailed Student t-test and all the data represented as n=3 and P value of p ≤ 0.05 was considered to be significant.

## Results

### Proteasomal inhibition facilitates expression of endodermal genes during spontaneous differentiation of mESCs

The UPS is known to play a pivotal role in regulating several cellular processes including pluripotency and differentiation but its role in cell fate determination remains an untapped area. Hence, we hypothesized our study to unravel the involvement of UPS in cell fate specification. As an initial step, we allowed mESCs to differentiate toward Embryoid bodies (EBs) and checked the efficiency of EB formation by transcript analysis (Suppl.Fig1b). We checked for the expression of three germ layer genes in the presence and absence of different concentrations (5nM & 10nM) of proteasome inhibitor (PI), MG132. The mRNA analysis of endoderm-specific genes *Sox17* and *Cxcr4* showed a 2.9 and 2-fold increase respectively upon 10nM PI treatment. In contrast, the expression of other representative lineage genes was either unmodulated or decreased (Fig 1a and c). At the protein level, we observed enhanced expression of the endodermal gene CXCR4, but not the ectodermal gene FOXG1 upon proteasomal inhibition (Fig 1b). At the concentration of PI used in the study, no alteration in KI67 staining was observed which signifies that the proliferation was unaffected (data not shown). Further, to corroborate the observation, we differentiated mESCs toward EB with and without 25µM proteasomal activator IU-1 and the transcript analysis demonstrated no increased expression of endoderm gene *Sox17*, whereas the mesodermal gene *Mixl1* and ectodermal genes *Foxg1* showed an increase in expression compared to undifferentiated mESCs (Fig 1d) and the protein level expression analysis by western blotting confirmed the same (Fig 1e). As the inhibition of proteasomal activity increased the endodermal gene expression, it was intriguing to analyze whether the proteasomal activity itself is modulated during the tri-lineage differentiation. mESCs were directed towards targeted tri-lineage differentiation with appropriate cytokine cues and assayed for proteasomal activity and observed a significant decrease in proteasomal activity in endodermal differentiated cells compared to other lineages and undifferentiated counterparts (Fig 1f). Based on the observation, we were interested in differentiating the mESCs to all three lineages in the presence and absence of MG132 and analyzing the expression of germ layer genes. Enhancement in the expression of endodermal genes *Sox17* and *Gata6*, but not in the expression of genes related to meso- and ecto-dermal layers were observed upon treatment with PI at transcript (Fig 1g) and protein level (Fig 1h). The above observation suggested proteasomal inhibition to favour endodermal differentiation.

**Figure 1:**
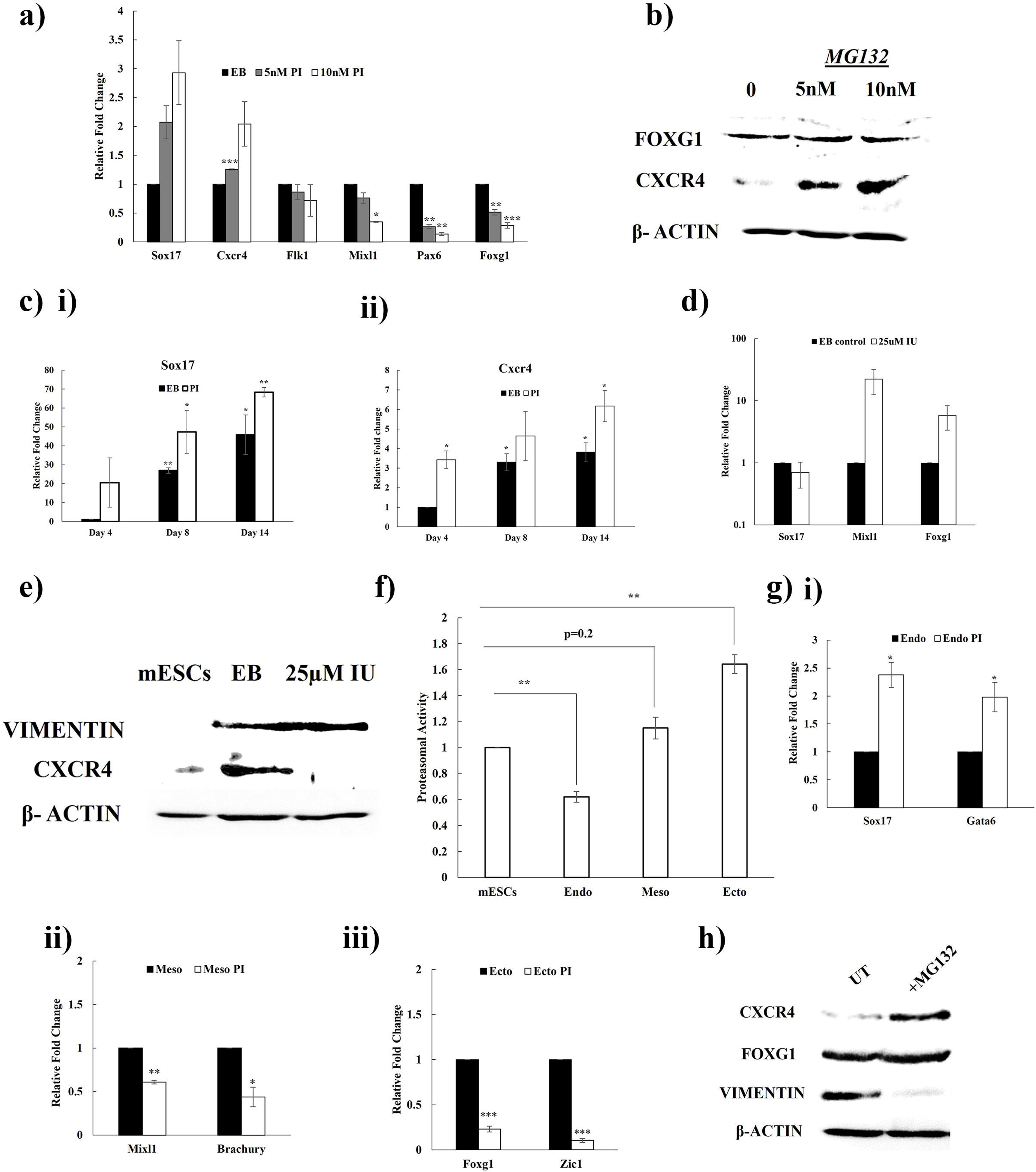
Enhanced expression of key endoderm-specific genes upon proteasome inhibition during spontaneous EB differentiation. a) Transcript analysis of the expression of tri-lineage markers during EB differentiation in presence and absence of different concentration of MG132, b) Immunoblot of EBs treated with 5 & 10nM PI for endoderm marker CXCR4 and ectoderm marker FOXG1, c) Gene expression level of endoderm markers (i) Sox17 and (ii) Cxcr4 during different day point of EB differentiation with 10nM PI, d) Relative gene expression analysis of representative tri-lineage genes during EB differentiation with and without IU-1, e) Western blot of EBs treated with 25µM IU-1 for endoderm and mesoderm lineage genes CXCR4 and Vimentin respectively, compared to undifferentiated mESCs, f) Chymo-trypsin like proteasomal activity in targeted tri-lineage differentiated cells compared to its undifferentiated counterparts, g) Transcript analysis of representative genes in targeted (i) endo-, (ii) meso-, (iii) ecto-derm differentiated cells in presence and absence of PI, h) Western blot analysis for the expression of representative genes of targeted trilineage differentiated cells cultured in presence or absence of PI. Data are represented as ±SE, (n=3); *, p ≤ 0.05, **, p ≤ 0.001, ***, p ≤ 0.001 compared to control.

### Proteasomal activity decreases during different stages of pancreatic β-cell differentiation of mESCs

To evaluate the effect of proteasomal inhibition during endodermal differentiation, we differentiated mESCs towards pancreatic β-cells with proper cytokine cues at regular stages of differentiation (Fig 2a). Transcript analysis at different day points revealed an increase in the expression of stage-specific genes compared to undifferentiated mESCs (Fig 2b). Gene expression of mesoderm markers Brachyury showed an increased expression in the early differentiation and later decreased with the day points (Fig 2c) and in contrast, expression of pluripotent gene *Oct4* was decreased during differentiation (Fig 2d), indicating the efficiency of the differentiation. Immunofluorescence staining of pancreatic progenitor marker HNF6 and β-cell mature marker Insulin demonstrated appropriate pancreatic islet cell differentiation (Fig 2e). In addition, we measured the number of C-peptide positive cells at the terminal stage of differentiation and observed 25.2%±0.5 positive cells in the islet-like cells compared to undifferentiated cells (Fig 2f). As experiments in this study suggested proteasomal inhibition to favour endodermal differentiation, we were interested in looking at the modulation in proteasomal activity during pancreatic differentiation. Analysis of proteasomal activity at different stages showed a gradual decrease in activity during differentiation (Fig 2g). In accordance, both embryonic and adult pancreas tissue showed lesser proteasomal activity compared to mESC control (Fig 2h). These results indicate the necessity of decreased proteasomal activity during pancreatic islet generation.

**Figure 2:**
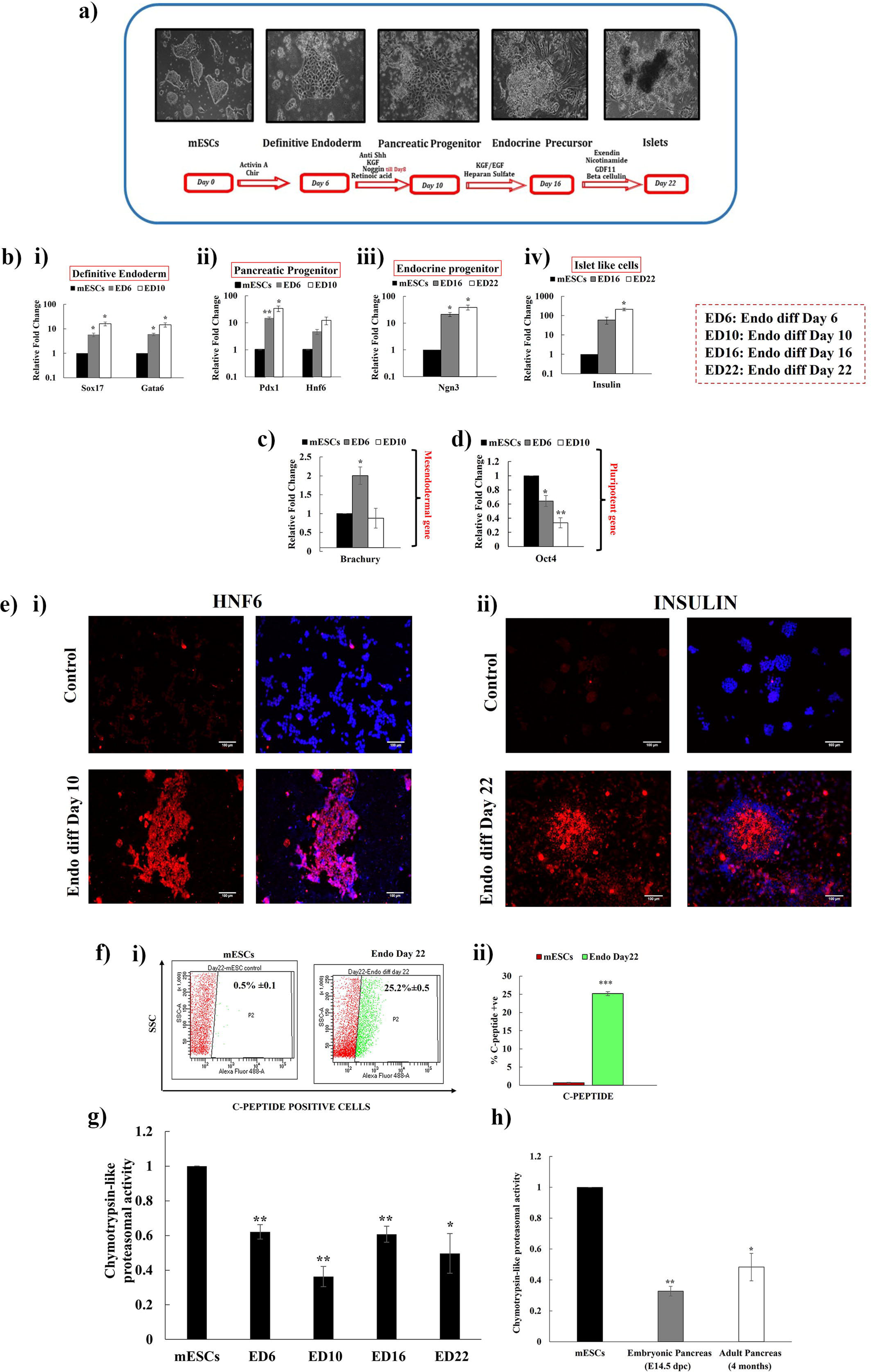
Decrease in Proteasomal activity during different stages of differentiation towards pancreatic β-cells from mESCs. a) Phase contrast representation stage-specific pancreatic β-cell differentiation, b) Relative gene expression level of representative genes of different stages of pancreatic β-cell differentiation compared to undifferentiated mESCs. c) Transcript analysis of mesendoderm gene *Brachury* and d) d) pluripotent gene *Oct4*, e) Immunofluorescence staining of (i)HNF6 and (ii) INS at differentiation day points 10 and 22 respectively (Scale bars=100µM), f) (i) Flow cytometry analysis for C-peptide positive cells in the terminally differentiated cells and (ii) quantified in comparison with undifferentiated mESCs, g) Analysis of chymo-trypsin like proteasomal activity during the course of endodermal differentiation and h) proteasomal activity analyzed in embryonic and adult pancreas compared to mESCs. Data are represented as ±SE, (n=3); *, p ≤ 0.05, **, p ≤ 0.001 compared to control.

### Inhibition of proteasomal activity enhances endodermal differentiation of mESCs

mESCs were differentiated towards pancreatic β-cells and by day 3 proteasomal inhibitor MG132 was supplemented with the differentiation cue till day 6 and the differentiated cells were harvested at different stages of pancreatic β-cell differentiation for analysis (Fig 3a). Gene expression analysis exhibited an increase in the expression of stage-specific genes during endodermal differentiation upon PI treatment compared to the untreated control (Fig 3b). The mesodermal gene Brachyury showed a decrease in the expression with proteasomal inhibition (Fig 3c). Protein analysis showed an upregulation of CXCR4 and PDX1 in the presence of MG132 compared to the control (Fig 3d). To determine whether the proteasomal activation reversed the effect, we induced endodermal differentiation in the presence and absence of proteasomal activator IU-1 and transcript analysis displayed a significant reduction in endodermal genes *Sox17* and *Gata6* (Fig 3e). Taken together, the data conclusively indicate proteasomal inhibition to enhance the endodermal differentiation of mESCs.

**Figure 3:**
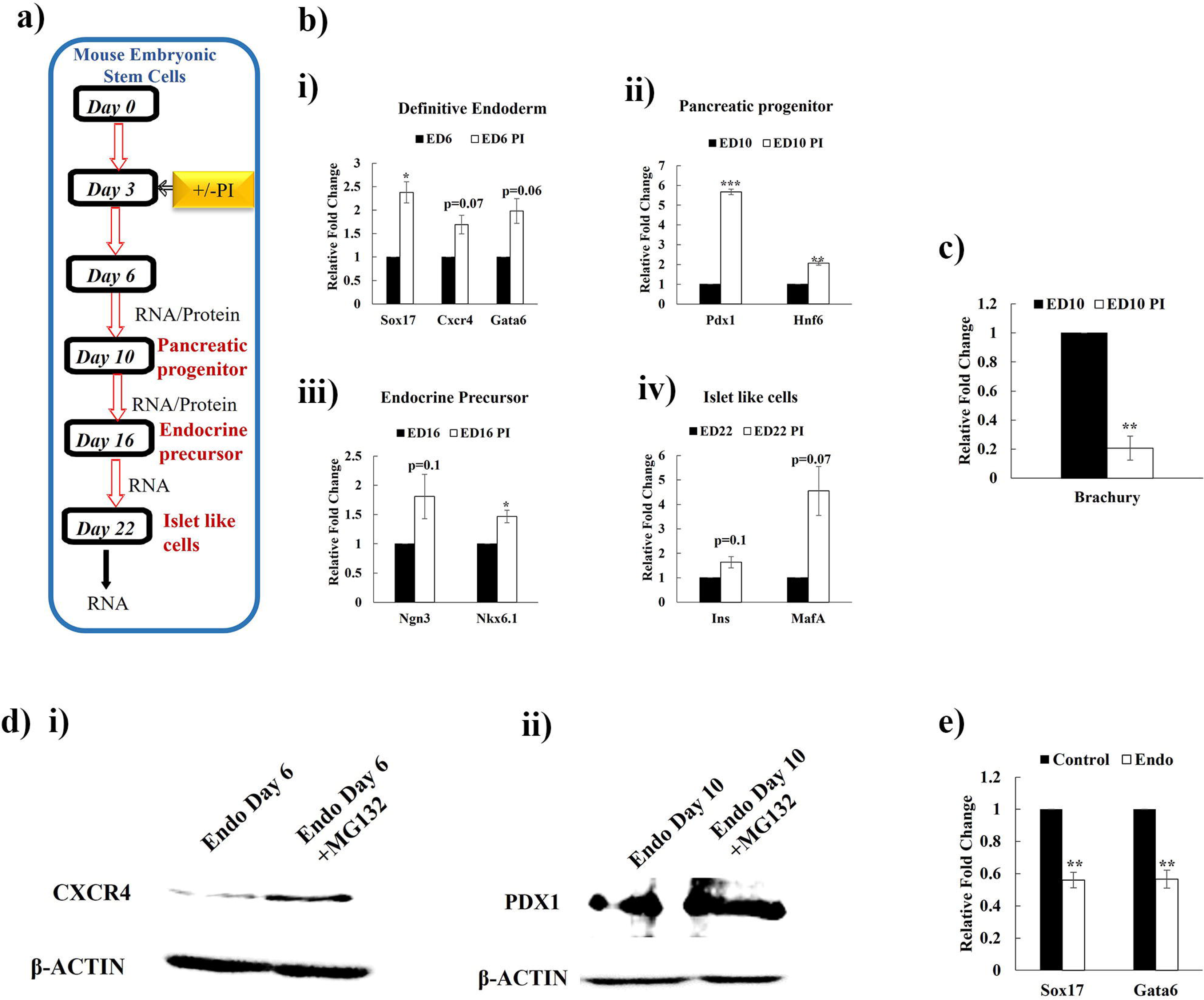
Inhibition of proteasomal activity enhances the β-cell differentiation of mESCs. a) Schematic representation of stage-specific differentiation of pluripotent stem cells to β-cells in the presence of proteasomal inhibitor, b) Transcript analysis of cells at different stages of (i) definitive endoderm, (ii) pancreatic progenitor, (iii) endocrine precursor, (iv) islet-like cells during the course of differentiation in presence or absence of PI c) mesendoderm marker Brachyury on day 10 of differentiation in presence or absence of PI d) Analysis of (i) CXCR4 and (ii) PDX1 expression at the protein level in presence or absence of PI in definitive endoderm differentiated cells, e) Relative gene expression analysis of endoderm core genes in presence or absence of proteasomal activator IU-1. Data are represented as ±SE, (n=3); *, p ≤ 0.05, **, p ≤ 0.001 compared to control.

### Inhibition of proteasomal activity enhances the generation of endoderm cells from mouse and human iPSCs

To confirm whether the observed effect of proteasomal inhibition on endodermal differentiation is restricted to mESCs, we used an independent reporter iPSC model system wherein expression of GFP is driven by PDX1 promoter (Pdx1-GFP iPSCs) (23). Pdx1-GFP iPSCs were differentiated to pancreatic β-cells in the presence and absence of MG132. Transcript analysis showed enhanced expression of stage-specific endodermal key genes in comparison to undifferentiated cells (Fig 4a). Flow cytometry analysis showed an enhanced number of GFP-positive cells in differentiated cells compared to undifferentiated cells. However, the cells treated with MG132 exhibited a greater number of GFP-positive cells compared to untreated cells (Fig 4b). We extended our observation to hiPSCs and proteasomal activity assay indicated a significant decrease in proteasomal activity in endodermal differentiated cells compared to undifferentiated hiPSCs (Fig 4c), as observed with mouse pluripotent stem cells (PSCs). This result confirmed the necessity of lower proteasomal activity during endodermal differentiation in both mouse and human PSCs. Inhibition of proteasomal activity during endodermal differentiation of hiPSCs exhibited enhanced expression of representative stage-specific genes compared to the untreated control (Fig 4d). Protein analysis of pancreatic progenitor gene PDX1 at different stages of endodermal differentiation showed increased expression in cells treated with MG132 compared to untreated cells (Fig 4e). These results conclusively demonstrated the importance of inhibition of proteasomal activity for enhanced endoderm differentiation in different PSCs model systems.

**Figure 4:**
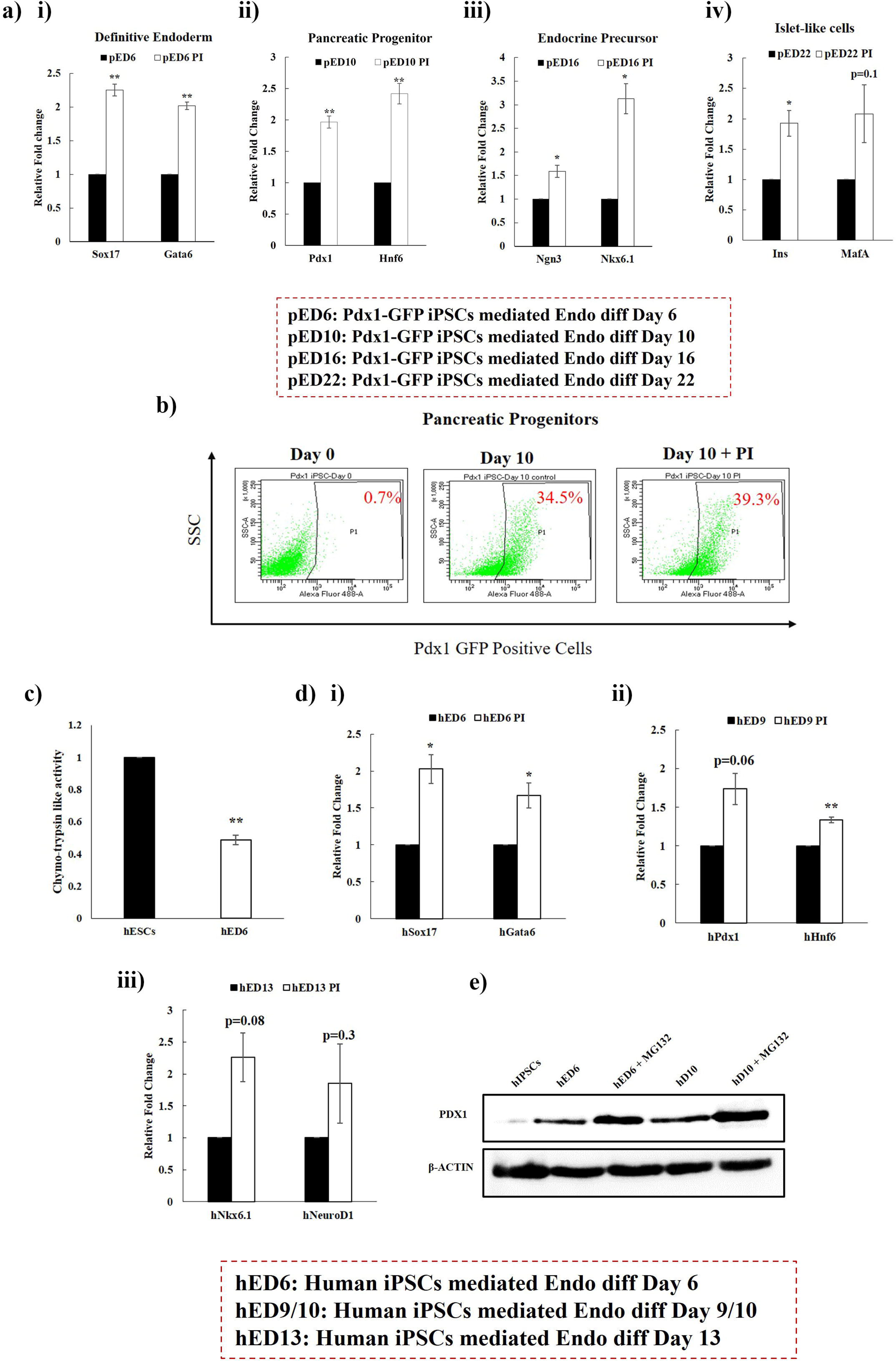
Proteasomal inhibition enhances the generation of endodermal cells from mouse and human iPSCs. a) Transcript analysis of cells from different stages of (i) definitive endoderm, (ii) pancreatic progenitor, (iii) endocrine precursor, (iv) and islet-like cells differentiated from Pdx1-GFP iPSCs in presence and absence of the proteasomal inhibitor, b) Flow cytometry assay for GFP positive cells from undifferentiated Pdx1-GFP iPSCs and cells differentiated to endoderm cells in presence or absence of PI. Data are represented as ±SE, (n=3) b, (n=1); *, p ≤ 0.05, **, p ≤ 0.001 compared to control. c) Chymo-trypsin-like proteasomal activity during hiPSCs-mediated endodermal differentiation, b) Real-time PCR analysis of endodermal genes at different day points during hiPSC differentiation in the presence and absence of PI c) Western blotting for PDX1 at different day points of hiPSC endoderm differentiation in presence or absence of PI. Data are represented as ±SE, (n=3); *, p ≤ 0.05, **, p ≤ 0.001 compared to control.

### Proteasome inhibition fails to elicit the Unfolded protein response during endodermal differentiation

To understand the mechanism behind enhanced endodermal differentiation upon proteasomal inhibition, we looked at the overall protein ubiquitination in the cells differentiated to trilineage. As expected, due to decreased proteasomal activity in endodermal lineage, we found an enhanced amount of ubiquitination in endodermal differentiation compared to other lineages (Fig 5a). To address the reason for the specific decrease in proteasomal activity in the endoderm lineage, we looked at the genes regulating the expression of proteasomal subunits and proteins involved in assembly. The expression of the regulatory genes was similar in all the tri-lineage differentiated cells, ruling out the possibility of their involvement in enhanced endodermal differentiation (Fig 5b). Previous studies showed increased ubiquitinated protein to activate the unfolded protein response (UPR). Studies also showed the UPR to be sufficient to generate the definitive endoderm specification from mESCs and found UPR inducers like Thapsigargin and tunicamycin to enhance the definitive endoderm cells (26). To detect the UPR activity in our study, we used a fluorogenic dye called TPE-MI (tetraphenylethene maleimide) which binds to the exposed cysteine upon denaturation and emits the fluorescence. mESCs treated in presence and absence of PI and thapsigargin showed an increase in the fluorescence in the cells treated with thapsigargin, but not with PI (Fig 5c). Targeted tri-lineage differentiated cells showed no difference in TPE-MI fluorescence intensity in all three germ layers (Fig 5d). Transcript analysis demonstrated no difference in the expression of UPR-related genes in the tri-lineage differentiated cells (Figure 5e). To validate the observation, the endoderm differentiation was performed in presence and absence of PI and TG and a significant increase in the expression of endoderm genes *Sox17* and *Gata6* was observed upon both PI and TG treatment (Figure 5f). To verify whether the effect of PI mediated enhancement in endodermal differentiation is via UPR, mESCs were allowed to differentiate towards endoderm in presence and absence of PI and TG. UPR genes *Atf4, Bip* and *Chop* showed elevated expression upon TG-mediated stress induction, but no modulation in expression was observed upon PI treatment in endoderm-differentiated cells (Figure 5g), indicating that enhanced endodermal differentiation upon PI is not due to a UPR response.

**Figure 5:**
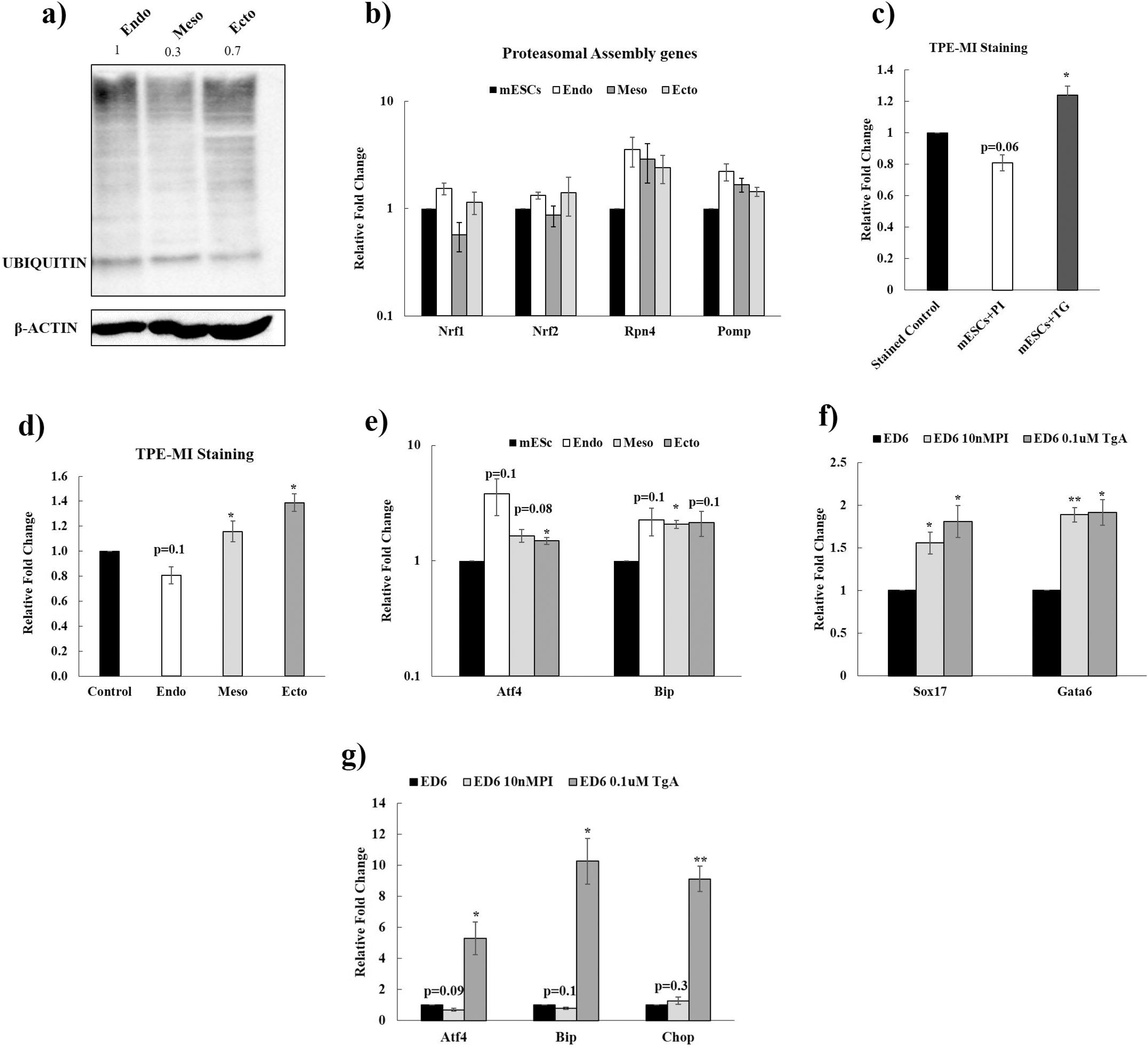
Enhanced endodermal differentiation upon proteasomal inhibition is not due to ER-UPR. a) Western blot assay for the ubiquitination pattern of total proteins in tri-lineage targeted differentiated cells, b) Gene expression analysis of proteasomal regulatory genes during targeted tri-lineage differentiation compared to its undifferentiated counterpart c) Unfolded protein load detection by TPE-MI staining on mESCs with and without PI and TG, d) TPE-MI staining for tri-lineage differentiated cells compared to mESCs, e) Transcript level of UPR genes by qPCR analysis in tri-lineage differentiated cells compared to mESCs, f) Real-time PCR analysis performed for the endodermal representative genes in presence or absence of PI and TG, g) Transcript level of UPR genes by qPCR analysis for the endodermal representative genes in presence or absence of PI and TG. Data is representative of ±SE, n=3 independent experiments (*p<0.05), (**p<0.01).

### The stability of YAP signalling mediates the enhanced endoderm differentiation by inhibition of proteasome

Inhibition of proteasomal activity during Days 6 and 10 of endodermal differentiation resulted in decreased expression of endodermal genes (Figure 6 a & b), whereas, as mentioned in the above results, the early inhibition (day 3 of differentiation) of proteasome enhanced the differentiation efficiency with increased expression of endodermal genes (Figure 6c). This observation inferred that early inhibition of proteasome disposes the cells towards endodermal differentiation. In the spree of identifying the mechanism, we further searched for the signaling pathways that influence early pancreatic development. The search identified the Hippo-Yap pathway as the promising candidate as the pathway is essential during the early stages of pancreatic differentiation and necessarily has to be downregulated at the later stages for efficient β-cell development (Table 1) (27). Previous reports have also demonstrated YAP1 to be the proteasomal substrate (28, 29). Hence, we were curious to see whether the YAP pathway is involved in the proteasome inhibition mediated enhanced endoderm differentiation. As an initial step, we evaluated the change in the expression pattern of *Yap* during endodermal differentiation and observed an upregulation of *Yap* expression in the early stage of differentiation and gradual reduction upon increasing day point (Figure 6 d & e). We looked at the expression of YAP during different day points of β-cell differentiation and observed an upregulation of YAP expression at Days 6 and 10 in the cells treated with PI, but during terminal differentiation of day 16 the YAP expression was undetected in both PI treated and untreated cells (Figure 6f). Additionally, we wanted to authenticate the role of YAP in the commitment of early endodermal differentiation. To this end, we used verteporfin-an inhibitor of the YAP/TEAD pathway, and subjected mESCs to endodermal differentiation. Activity of verteporfin was confirmed by the downregulation in the expression of *Gli2*, the downstream target of YAP (Figure 6g). Further analysis revealed a drastic reduction in the expression of multipotent pancreatic progenitor markers *Sox9, Notch1,* and *Smad6* upon treatment with YAP inhibitor verteporfin compared to differentiated cells. Interestingly, the reduced expression due to verteporfin was restored upon proteasomal inhibition (Figure 6 h & i), indicating the involvement of the YAP pathway in proteasome inhibition-mediated increase in endodermal differentiation.

**Figure 6:**
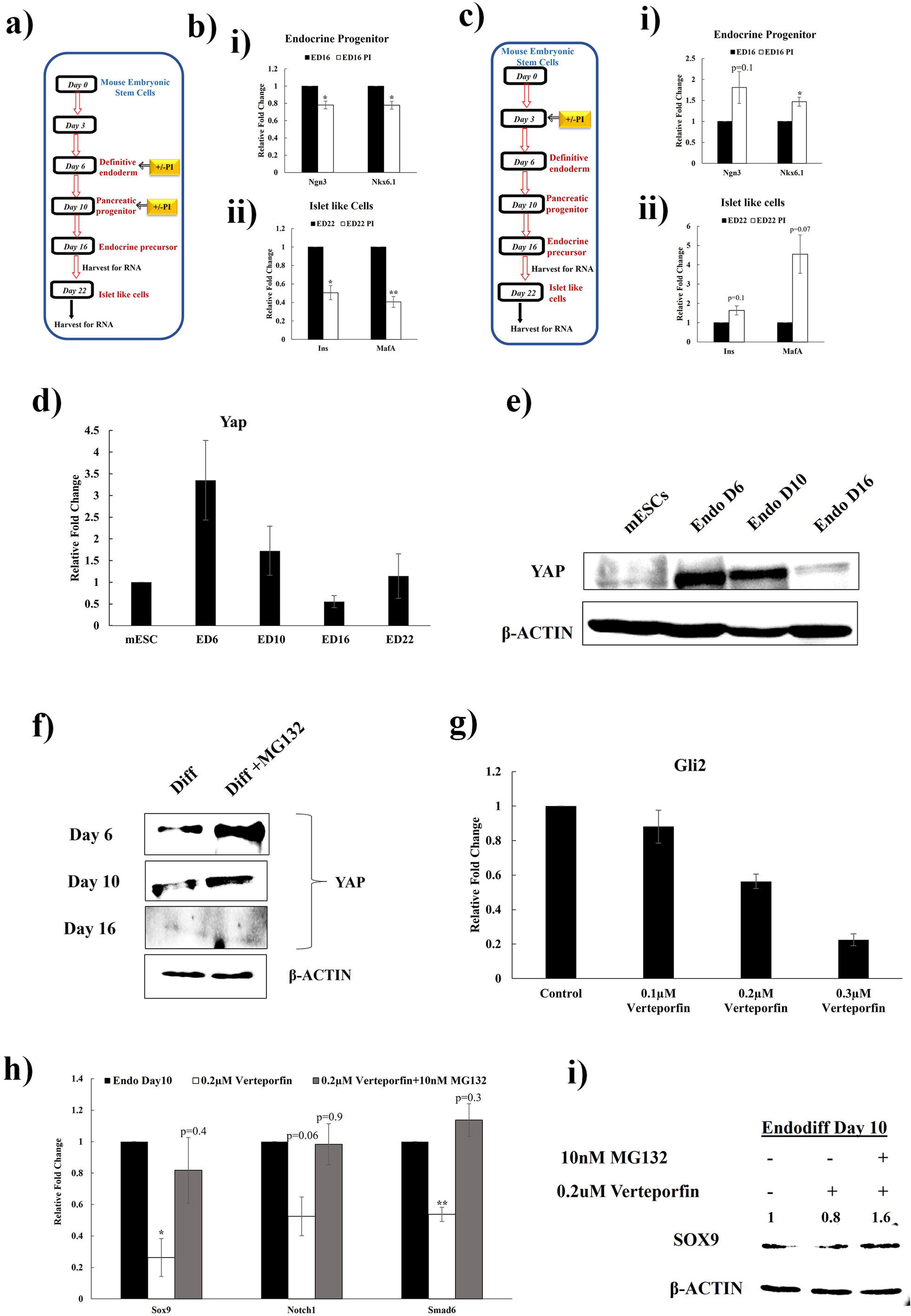
Inhibition of proteasome enhances the stability of YAP resulting in enhanced endoderm differentiation: a) Schematic representation of differentiation of mESCs to islet-like cells and addition of PI and different stages of differentiation. b) Gene expression analysis of (i) endocrine progenitor and (ii) matured islet genes in cells differentiated with the addition of PI either on Day 6 or Day 10. c) Schematic representation and gene expression analysis of (i) endocrine progenitor and (ii) matured islet genes in cells differentiated with the addition of PI on Day 3 of differentiation. Transcript (d) and protein analysis (e) of YAP during different time points of endoderm differentiation. f) Analyzing the stability of YAP at the protein level during differentiation in cells treated with or without PI. g) Transcript level of YAP downstream target gene in cells cultured with different concentrations of YAP inhibitor Verteporfin. h) Transcript and i) protein analysis of representative genes of pancreatic multipotent progenitor genes in cells cultured in the presence of verteporfin and treated with or without PI. Data is representative of ±SE, n=3 independent experiments (*p<0.05), (**p<0.01).

**Table 1:**
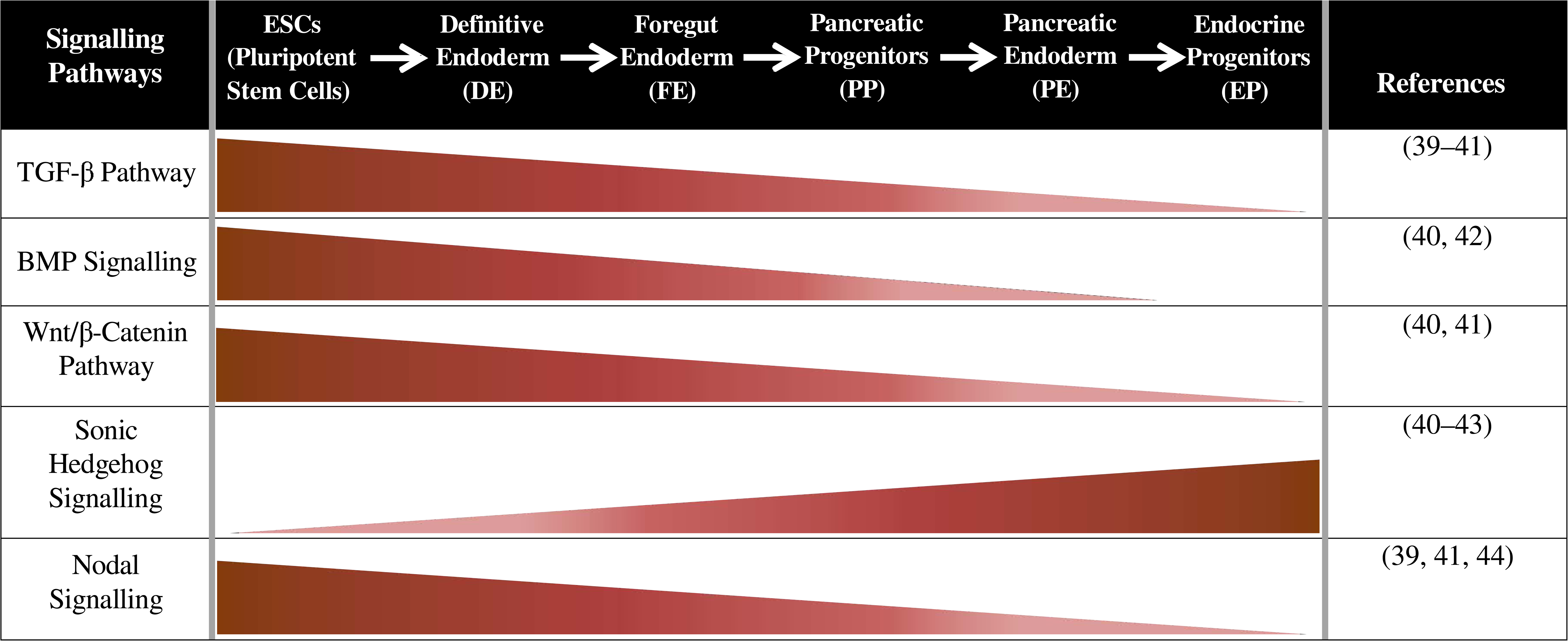
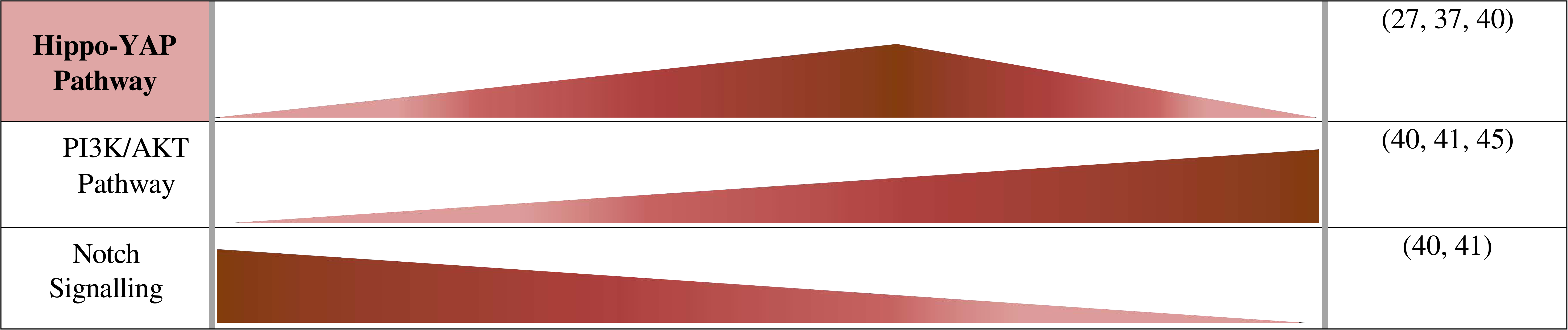
Stage specific involvement of different key signaling pathways during Pancreatic β-cell development.

## Discussion

The fidelity of the pluripotent inner cell mass to differentiate into cells of different germ layers is largely dependent on accurate proteostasis maintained by selective proteolytic mechanism (30). Previous studies demonstrated pluripotent stem cells to have increased proteasomal activity compared to their differentiated counterparts (13). Inhibition of proteasomal activity induced the demise of pluripotent identity with concomitant gain in the expression of differentiation genes (13, 31). UPS plays a significant role in maintaining the homeostasis of cellular proteins which is pivotal for the inner cell mass to decide the fate of the cells during development. During pancreatic development, several key proteins that regulate different stages of development are substrates of proteasome. Though many of the TFs that participate in pancreatic development have been shown to be substrates of proteasomes in different tissues, their direct involvement in pancreatic tissue development is relatively less understood (32). For instance, the homeostasis of TF CXCR4 is regulated by E3 ligase AIP4 and deubiquitinase USP14 (33, 34). However, its role has been revealed in cancer cells and whether similar enzymes participate in the pancreas is yet to be revealed (34). The turnover of few TFs such as PDX1 and NGN3 by proteasome has been revealed in the pancreas itself. FBXW7, an E3 ubiquitin ligase belonging to the SCF (Complex of SKP1, CUL1, and F-box protein) family ubiquitinates NGN3 protein and relays it to proteasomal degradation (9, 35). The involvement of proteasomes in pancreatic development intrigued us to investigate whether inhibition of proteasomal activity in pluripotent stem cells (PSCs) has a preferential proclivity to differentiate into one of the three germ layers. In this study, we show that inhibition of proteasomal activity coaxes ESCs towards endoderm lineage.

To the best of our knowledge, this is the first report that shows UPS, despite having a pleiotropic cellular function, can facilitate definitive endodermal differentiation in *in vitro* systems of different culture conditions like EBs and targeted differentiation of PSCs. Inhibition of proteasomal chymotryptic activity by MG132 either during EB formation or targeted differentiation enhanced the generation of definitive endodermal lineage cells from mouse ESCs, iPSCs as well from human iPSCs without altering cell proliferation (data not shown). Supplementing the proteasomal activity activator IU1 decreased the endodermal commitment and concomitantly enhanced the meso- and ecto-dermal genes. A decrease in proteasomal activity during differentiation and ageing has been well-documented, but the inhibition of proteasomal activity to have a larger impact on endodermal differentiation is the novel outcome of this study. These observations probably emphasize the prerequisite of declining the proteasomal activity for definitive endodermal cell fate decision during gastrulation process.

To understand the underlying mechanism, we started our investigation focusing on two aspects: a) The reason behind the specific decrease in proteasomal activity in endoderm differentiation and b) how a decrease in proteasomal activity results in enhanced endodermal differentiation. To address the first query, we analyzed the expression level of the known regulators of the expression of proteasomal subunits. However, there was no detectable difference in the expression of the regulators. Previous studies by Xu et al., had reported that the unfolded protein response (UPR) activity is required for endoderm specification, and this is achieved by modulating the WNT and TGFβ signaling pathways (26). Also, an independent study showed the inhibition of proteasomal activity to enhance UPR (36), and therefore we hypothesized UPR to be an intermediate player and sought to delineate the role of UPR behind the enhanced endoderm specification by inhibiting proteasomal activity. However, no detectable differences were identified in the expression of UPR genes in the presence or absence of PI and the expression was similar in all three lineages, thereby ruling out the contribution of UPR in proteasome inhibition-mediated enhanced endoderm differentiation. In this study, we failed to address the reason behind the decrease in proteasomal activity specifically in the endoderm lineage.

To address the second aspect of how a decrease in proteasomal activity results in enhanced endodermal differentiation, we analyzed the effect of temporal inhibition of proteasome during differentiation of ESCs and found enhanced endodermal differentiation only when proteasome was inhibited at an early stage of differentiation. Screening for different signaling pathways that have a profound role during early differentiation of endoderm, identified the YAP-signaling pathway to be the candidate. YAP has been shown to play a significant role in multipotent progenitor generation by regulating enhancer elements of transcription factors involved in the processes (37). However, for terminal differentiation, YAP is supposed to undergo phosphorylation and further proteasome-mediated degradation (38). Downregulation of YAP has been documented to be necessary for the cells to transit from the multipotent progenitor stage to NGN3+ endocrine progenitors and islet cells (27). Expression analysis in our study also showed an upregulation of YAP expression at both transcript and protein levels during the early stages of endoderm differentiation and gradual reduction upon increasing day point. Previous studies have documented the YAP protein to be the substrate of proteasome (28). The YAP TF is phosphorylated by Lats and CK1 delta/epsilon in a phosphodegron and recruits the SCF (beta-TRCP) E3 ubiquitin ligase, which ubiquitinates YAP and channelizes to proteasome-mediated degradation (29). In the present study, the result indicated that the addition of PI enhanced the stability of the YAP protein and facilitated the increased generation of pancreatic multipotent progenitor cells. Also, we observed partial rescue in the expression of multipotent progenitor genes when simultaneously YAP signaling is inhibited by verteporfin and proteasome inhibited by PI. Probably, stabilizing YAP at the early stage of differentiation by inhibiting proteasome could be one of the mechanisms by which inhibition of proteasome at the early stage of differentiation facilitated enhanced endodermal differentiation from PSCs.

In summary, our study highlights the importance of decreasing proteasomal activity to achieve better endodermal differentiation from PSCs and the attempt to address the mechanism sheds light on the YAP signaling pathway as one of the underlying mechanisms. Further studies will be needed to establish the existence of alternative mechanisms as the simultaneous presence of YAP inhibitor verteporfin and PI rescued the expression of multipotent progenitor genes partially. In conclusion, the study highlights the specific role of proteasome during germ layer specification and this study accentuates the possible application of PI as an ingredient in the cocktail of cytokines to drive the PSCs to endoderm cells.

## Acknowledgement

This study was supported by grants from the Science and Engineering Research Board Department of Science and Technology, Government of India (CRG/2021/000960) to AK. PDX-GFP iPSC is a kind gift from Prof. Catherine Verfaillie, katholieke universiteit, Leuven. We thank AK lab members for their technical help, suggestions and encouragement.

## Author contributions

Conceptualization, A.A., J.P., and A.K.; Methodology and Formal Analysis, A.A., S.S., and A.K.; Investigation, A.A., S.S., and A.K.; Writing – Original Draft, A.A., and A.K.; Writing – Review & Editing: A.A., S.S., J.P., and A.K.; Supervision, A.K.; Funding Acquisition, A.K.

## Declaration of interests

The authors declare no competing interest.

## Data and Code Availability

The data that support the findings of this study are available from the Lead Contact on reasonable request.

## Notes

### Competing Interest Statement

The authors have declared no competing interest.

## Reference

1. Abdul Basith Khan, M., Hashim, M. J., King, J. K., Govender, R. D., Mustafa, H., and Al Kaabi, J. (2020) Epidemiology of type 2 diabetes—global burden of disease and forecasted trends. J. Epidemiol. Glob. Health. 10, 107–111

2. Soumya, D., and Srilatha, B. (2011) Late stage complications of diabetes and insulin resistance. J Diabetes Metab. 2, 1000167

3. Pecoits-Filho, R., Abensur, H., Betonico, C. C. R., Machado, A. D., Parente, E. B., Queiroz, M., Salles, J. E. N., Titan, S., and Vencio, S. (2016) Interactions between kidney disease and diabetes: dangerous liaisons. Diabetol. Metab. Syndr. 8, 1–21

4. Neumann, M., Arnould, T., and Su, B.-L. (2023) Encapsulation of stem-cell derived β-cells: A promising approach for the treatment for type 1 diabetes mellitus. J. Colloid Interface Sci. 636, 90–102

5. Song, J., and Millman, J. R. (2016) Economic 3D-printing approach for transplantation of human stem cell-derived β-like cells. Biofabrication. 9, 15002

6. Murtaugh, L. C., and Melton, D. A. (2003) Genes, signals, and lineages in pancreas development. Annu. Rev. Cell Dev. Biol. 19, 71–89

7. Pagliuca, F. W., and Melton, D. A. (2013) How to make a functional β-cell. Development. 140, 2472–2483

8. Klein, S., Meng, R., Montenarh, M., and Götz, C. (2016) The phosphorylation of PDX-1 by protein kinase CK2 is crucial for its stability. Pharmaceuticals. 10, 2

9. Sancho, R., Gruber, R., Gu, G., and Behrens, A. (2014) Loss of Fbw7 Reprograms Adult Pancreatic Ductal Cells into α, δ, and β Cells. Cell Stem Cell. 10.1016/j.stem.2014.06.019

10. Wang, J., and Maldonado, M. A. (2006) The ubiquitin-proteasome system and its role in inflammatory and autoimmune diseases. Cell Mol Immunol. 3, 255–261

11. Saric, T., Graef, C. I., and Goldberg, A. L. (2004) Pathway for degradation of peptides generated by proteasomes: a key role for thimet oligopeptidase and other metallopeptidases. J. Biol. Chem. 279, 46723–46732

12. Nandi, D., Tahiliani, P., Kumar, A., and Chandu, D. (2006) The ubiquitin-proteasome system. J. Biosci. 10.1007/BF02705243

13. Vilchez, D., Boyer, L., Morantte, I., Lutz, M., Merkwirth, C., Joyce, D., Spencer, B., Page, L., Masliah, E., and Berggren, W. T. (2012) Increased proteasome activity determines human embryonic stem cell identity. Nature. 489, 304

14. Saez, I., Koyuncu, S., Gutierrez-Garcia, R., Dieterich, C., and Vilchez, D. (2018) Insights into the ubiquitin-proteasome system of human embryonic stem cells. Sci. Rep. 8, 4092

15. Hartley, T., Brumell, J., and Volchuk, A. (2009) Emerging roles for the ubiquitin-proteasome system and autophagy in pancreatic β-cells. Am. J. Physiol. Metab. 296, E1– E10

16. Boucher, M.-J., Selander, L., Carlsson, L., and Edlund, H. (2006) Phosphorylation marks IPF1/PDX1 protein for degradation by glycogen synthase kinase 3-dependent mechanisms. J. Biol. Chem. 281, 6395–6403

17. Fujimoto, K., Ford, E. L., Tran, H., Wice, B. M., Crosby, S. D., Dorn, G. W., and Polonsky, K. S. (2010) Loss of Nix in Pdx1-deficient mice prevents apoptotic and necrotic β cell death and diabetes. J. Clin. Invest. 120, 4031–4039

18. Liang, J., Chirikjian, M., Pajvani, U. B., and Bartolomé, A. (2022) MafA regulation in β-cells: from transcriptional to post-translational mechanisms. Biomolecules. 12, 535

19. Wu, T., Zhang, S., Xu, J., Zhang, Y., Sun, T., Shao, Y., Wang, J., Tang, W., Chen, F., and Han, X. (2020) HRD1, an important player in pancreatic β-cell failure and therapeutic target for type 2 diabetic mice. Diabetes. 69, 940–953

20. Kanai, K., Aramata, S., Katakami, S., Yasuda, K., and Kataoka, K. (2011) Proteasome activator PA28 stimulates degradation of GSK3-phosphorylated insulin transcription activator MAFA. J. Mol. Endocrinol. 47, 119–127

21. Kawaguchi, M., Minami, K., Nagashima, K., and Seino, S. (2006) Essential Role of Ubiquitin-Proteasome System in Normal Regulation of Insulin Secretion. J. Biol. Chem. 10.1074/jbc.M601228200

22. López-Avalos, M. D., Duvivier-Kali, V. F., Xu, G., Bonner-Weir, S., Sharma, A., and Weir, G. C. (2006) Evidence for a role of the ubiquitin-proteasome pathway in pancreatic islets. Diabetes. 55, 1223–1231

23. Porciuncula, A., Kumar, A., Rodriguez, S., Atari, M., Araña, M., Martin, F., Soria, B., Prosper, F., Verfaillie, C., and Barajas, M. (2016) Pancreatic differentiation of Pdx1-GFP reporter mouse induced pluripotent stem cells. Differentiation. 92, 249–256

24. Pagliuca, F. W., Millman, J. R., Gürtler, M., Segel, M., Van Dervort, A., Ryu, J. H., Peterson, Q. P., Greiner, D., and Melton, D. A. (2014) Generation of Functional Human Pancreatic β Cells In Vitro. Cell. 10.1016/j.cell.2014.09.040

25. Kisselev, A. F., and Goldberg, A. L. (2005) Monitoring activity and inhibition of 26S proteasomes with fluorogenic peptide substrates. Methods Enzymol. 398, 364–378

26. Xu, H., Tsang, K. S., Wang, Y., Chan, J. C. N., Xu, G., and Gao, W.-Q. (2014) Unfolded protein response is required for the definitive endodermal specification of mouse embryonic stem cells via Smad2 and β-catenin signaling. J. Biol. Chem. 289, 26290– 26301

27. Rosado-Olivieri, E. A., Anderson, K., Kenty, J. H., and Melton, D. A. (2019) YAP inhibition enhances the differentiation of functional stem cell-derived insulin-producing β cells. Nat. Commun. 10, 1464

28. Yan, C., Yang, H., Su, P., Li, X., Li, Z., Wang, D., Zang, Y., Wang, T., Liu, Z., and Bao, Z. (2022) OTUB1 suppresses Hippo signaling via modulating YAP protein in gastric cancer. Oncogene. 41, 5186–5198

29. Zhao, B., Li, L., Tumaneng, K., Wang, C.-Y., and Guan, K.-L. (2010) A coordinated phosphorylation by Lats and CK1 regulates YAP stability through SCFβ-TRCP. Genes Dev. 24, 72–85

30. Yan, P., Ren, J., Zhang, W., Qu, J., and Liu, G.-H. (2020) Protein quality control of cell stemness. Cell Regen. 9, 1–11

31. Atkinson, S. P., Collin, J., Irina, N., Anyfantis, G., Kyung, B. K., Lako, M., and Armstrong, L. (2012) A putative role for the immunoproteasome in the maintenance of pluripotency in human embryonic stem cells. Stem Cells. 30, 1373–1384

32. Aoki, M., Jiang, H., and Vogt, P. K. (2004) Proteasomal degradation of the FoxO1 transcriptional regulator in cells transformed by the P3k and Akt oncoproteins. Proc. Natl. Acad. Sci. 101, 13613–13617

33. Bhandari, D., Robia, S. L., and Marchese, A. (2009) The E3 ubiquitin ligase atrophin interacting protein 4 binds directly to the chemokine receptor CXCR4 via a novel WW domain-mediated interaction. Mol. Biol. Cell. 20, 1324–1339

34. Mines, M. A., Goodwin, J. S., Limbird, L. E., Cui, F.-F., and Fan, G.-H. (2009) Deubiquitination of CXCR4 by USP14 is critical for both CXCL12-induced CXCR4 degradation and chemotaxis but not ERK activation. J. Biol. Chem. 284, 5742–5752

35. Seifert, B. A., and Xiong, Y. (2014) Out of the F-box: reawakening the pancreas. Cell Stem Cell. 15, 111–112

36. Obeng, E. A., Carlson, L. M., Gutman, D. M., Harrington Jr, W. J., Lee, K. P., and Boise, L. H. (2006) Proteasome inhibitors induce a terminal unfolded protein response in multiple myeloma cells. Blood. 107, 4907–4916

37. Cebola, I., Rodríguez-Seguí, S. A., Cho, C. H. H., Bessa, J., Rovira, M., Luengo, M., Chhatriwala, M., Berry, A., Ponsa-Cobas, J., and Maestro, M. A. (2015) TEAD and YAP regulate the enhancer network of human embryonic pancreatic progenitors. Nat. Cell Biol. 17, 615–626

38. Ardestani, A., and Maedler, K. (2018) The Hippo signaling pathway in pancreatic β-cells: Functions and regulations. Endocr. Rev. 39, 21–35

39. Kim, S. K., and Hebrok, M. (2001) Intercellular signals regulating pancreas development and function. Genes Dev. 15, 111–127

40. Jin, W., and Jiang, W. (2022) Stepwise differentiation of functional pancreatic β cells from human pluripotent stem cells. Cell Regen. 11, 24

41. Tsaniras, C., and Jones, P. M. (2010) Generating pancreatic beta-cells from embryonic stem cells by manipulating signaling pathways. J. Endocrinol. 206, 13–26

42. Kumar, S. S., Alarfaj, A. A., Munusamy, M. A., Singh, A. J. A. R., Peng, I.-C., Priya, S. P., Hamat, R. A., and Higuchi, A. (2014) Recent developments in β-cell differentiation of pluripotent stem cells induced by small and large molecules. Int. J. Mol. Sci. 15, 23418– 23447

43. Hawkins, K., Joy, S., and McKay, T. (2014) Cell signalling pathways underlying induced pluripotent stem cell reprogramming. World J. Stem Cells. 6, 620

44. Sulzbacher, S., Schroeder, I. S., Truong, T. T., and Wobus, A. M. (2009) Activin A-induced differentiation of embryonic stem cells into endoderm and pancreatic progenitors—the influence of differentiation factors and culture conditions. Stem Cell Rev. Reports. 5, 159–173

45. Huang, H., Bader, T. N., and Jin, S. (2020) Signaling molecules regulating pancreatic endocrine development from pluripotent stem cell differentiation. Int. J. Mol. Sci. 21, 5867

