## Supplemental Table for "Inhibition of Proteasome Activity Facilitates Definitive Endodermal Specification of pluripotent Stem Cells by influencing YAP signaling"

**Supplementary Table 1:** The list of primer sequences of genes used in this study.

| No. | **Genes** | **Sequences** |
| --- | --- | --- |
| 1 | *mGapdh F* | ACCACAGTCCATGCCATCAC |
|  | *mGapdh R* | TCCACCACCCTGTTGCTGTA |
| 2 | *mSox17 F* | GATGCGGGATACGCCAGTG |
|  | *mSox17 R* | CCACCACCTCGCCTTTCAC |
| 3 | *mCxcr4 F* | CTTCTGGGCAGTTGATGCCAT |
|  | *mCxcr4 R* | CTGTTGGTGGCGTGGACAAT |
| 4 | *mFlk1 F* | CCCGCATGAAATTGAGCTAT |
|  | *mFlk1 R* | AAACATCTTCGCCACAGTCC |
| 5 | *mMixl1 F* | ACGCAGTGCTTTCCAAACC |
|  | *mMixl1 R* | CCCGCAAGTGGATGTCTGG |
| 6 | *mPax6 F* | CGGCAGAAGATCGTAGAG |
|  | *mPax6 R* | GATGACACACTGGGTATG |
| 7 | *mFoxg1 F* | CTGAGTGTGGACCGGCTG |
|  | *mFoxg1 R* | CGTGCTGGTCTGCGAAGTC |
| 8 | *mGata6 F* | CAACACAGTCCCCGTTCTTT |
|  | *mGata6 R* | TGGTACAGGCGTCAAGAGTG |
| 9 | *mBrachyury F* | TCCCGAGACCCAGTTCATAG |
|  | *mBrachyury R* | TTCTTTGGCATCAAGGAAGG |
| 10 | *mZic1 F* | GCCCTTCAAAGCCAAATACA |
|  | *mZic1 R* | TTGCAAAGGTAGGGCTTGTC |
| 11 | *mPdx1 F* | ACAGCCCTGAGCTTCTGAAA |
|  | *mPdx1 R* | CTGCTGGTCCGTATTGGAAC |
| 12 | *mHnf6 F* | CCCTGGAGCAAACTCAAGTC |
|  | *mHnf6 R* | GCTGGGAGATGGTGATTTGT |
| 13 | *mNgn3 F* | GAGTTGGCACTCAGCAAACA |
|  | *mNgn3 R* | TCTGAGTCAGTGCCCAGATG |
| 14 | *mNkx6.1 F* | TCAGGTCAAGGTCTGGTTCC |
|  | *mNkx6.1 R* | CGATTTGTGCTTTTTCAGCA |
| 15 | *mIns1 F* | TGTTGGTGCACTTCCTACCC |
|  | *mIns1 R* | GCTGGTAGAGGGAGCAGATG |
| 16 | *mMafA F* | CGGGACCTGTACAAGGAGAA |
|  | *mMafA R* | GGCACGTCACAGAAAGAAGTC |
| 17 | *mOct4 F* | CTGCTGAAGCAGAAGAGGATCAC |
|  | *mOct4 R* | CTTCTGGCGCCGGTTACAGAACCA |
| 18 | *mNrf1F* | CCCCCGAGGACACTTCTTAT |
|  | *mNrf1R* | CGTGTCTGCTGTCTCTTTCG |
| 19 | *mNrf2 F* | GGTTGCCCACATTCCCAAAC |
|  | *mNrf2 F* | ACTTGCTCCATGTCCTGCTC |
| 20 | *mRpn4 F* | TCACAAGGCACTGATGAAGC |
|  | *mRpn4 R* | CGGGGCTGATACTGTTCACT |
| 21 | *mPomp F* | TCTCCGGAAAGGGTTTTCTT |
|  | *mPomp R* | CTTCAGGGGAGCAAACAGAC |
| 22 | *mAtf4 F* | TTAGAGCTAGGCAGTGAAGTT |
|  | *mAtf4 R* | CTGTCATTGTCAGAGGGAGT |
| 23 | *mBip F* | CCATCCCGTGGCATAAAC |
|  | *mBip R* | GACTCCTCCCACAGTTTCA |
| 24 | *mChop F* | CACATCCCAAAGCCCTCG |
|  | *mChop R* | CGTTCTCCTGCTCCTTCTC |
| 25 | *mYap F* | ACCCTCGTTTTGCCATGAAC |
|  | *mYap R* | CCTTCTCCATCTGTAACTGC |
| 26 | *mGli2 F* | GGTCAAGACTGAGGCTGAGG |
|  | *mGli2 R* | CAGCTGCTCCTGTGTGTCAT |
| 27 | *mSox9 F* | CACACAGCTCACTCGACCTT |
|  | *mSox9 R* | AAGTGGGTAATGCGCTTGGA |
| 28 | *mSmad6 F* | AGTGGAGCTGAAACCCCTGT |
|  | *mSmad6 R* | AGGAGGAGACAGCCGAGAAT |
| 29 | *mNotch1 F* | GAGATGCTCCCAGCCAAGT |
|  | *mNotch1 R* | TCTTACACGGTGTGCTGAGG |
| 30 | *hGAGDH F* | TGGTATCGTGGAAGGACTCATGAC |
|  | *hGAGDH R* | ATGCCAGTGAGCTTCCCGTTCAGC |
| 31 | *hSOX17 F* | CGCACGGAATTTGAACAGTA |
|  | *hSOX17 R* | GGATCAGGGACCTGTCACAC |
| 32 | *hGATA6 F* | TCTCCATGTGCATTGGGGAC |
|  | *hGATA6 R* | AAGGAAATCGCCCTGTTCGT |
| 33 | *hPDX1 F* | ACCAAAGCTCACGCGTGGAAA |
|  | *hPDX1 R* | TGATGTGTCTCTCGGTCAAGTT |
| 34 | *hHNF6 F* | CGCTCCGCTTAGCAGCAT |
|  | *hHNF6 R* | GTGTTGCCTCTATCCTTCCCAT |
| 35 | *hNKX6.1F* | CTGGCCTGTACCCCTCATCA |
|  | *hNKX6.1R* | CTTCCCGTCTTTGTCCAACAA |
| 36 | *hNEUROD1 F* | GGATGACGATCAAAAGCCCAA |
|  | *hNEUROD1 R* | GCGTCTTAGAATAGCAAGGCA |
